# Spatial organization of self-assembled G-quadruplex nucleic acids tunes extracellular electron transfer in pathogenic biofilms

**DOI:** 10.64898/2026.06.17.732861

**Authors:** Obinna M. Ajunwa, Rikke L. Meyer

## Abstract

Extracellular electron transfer (EET) allows bacteria to sustain metabolism when soluble electron acceptors such as oxygen are scarce, a situation typical of the biofilm interior. These mechanisms are well characterised in environmental metal-reducing bacteria but poorly defined in the biofilms of pathogens, where they may contribute to persistence at infection sites. We previously found that synthetic guanine-quadruplex (G4) nucleic acids bound to hemin can conduct electrons^1^, yet whether bacteria self-assemble such structures in electroactive form was unknown.

Here we show that *Staphylococcus aureus* biofilms naturally assemble G4-rich extracellular nucleic acids that bind hemin to form a catalytically active, electron-conducting complex, without exogenous G4 addition. Nutrient starvation, rather than biofilm age or cell density, triggers extracellular G4 accumulation, and as biofilms develop, the G4 reorganise from intercellular networks to being primarily located at the cell-envelope. Using electrochemistry, peroxidase imaging and different types of nucleases, we show that these architectures impose distinct modes of electron transfer through the matrix or at the cell surface. Degradation of G4 abolishes electroactivity whereas removing canonical B-DNA does not. *S. aureus* thus builds and organises its own electroactive nucleic-acid network, identifying biofilm architecture as a tunable determinant of extracellular electron flow.

## Introduction

Extracellular electron transfer (EET) allows bacteria to move electrons between their intracellular metabolism and the external environment, sustaining energy conservation when soluble terminal electron acceptors such are scarce^2,3^. This is the case in biofilms that contain steep chemical gradients, and here EET supports metabolic activity and intercellular interaction^4,5^. Mechanisms for EET are well characterised in environmental metal-reducers such as *Geobacter* and *Shewanella*, which deploy multiheme cytochromes, conductive appendages, and secreted redox shuttles^6^. By contrast, whether and how EET operates in the biofilms of bacterial pathogens remain poorly understood, despite its potential for impacting persistence and antimicrobial tolerance in biofilm infections.

We recently described an unconventional EET pathway that does not depend on dedicated redox proteins: guanine-quadruplex (G4) nucleic acids that bind hemin (Fe³⁺-protoporphyrin IX) to form a redox-active complex capable of conducting electrons in *Staphylococcus epidermidis* biofilms^1^. G4s are non-canonical nucleic acid structures of stacked guanine tetrads from one or multiple strands held together by Hoogsteen hydrogen bonds^7^, and the G4/hemin complex behaves as an oxidoreductase^8,9^. A conceptually similar but hemin-free G4-RNA EET conduit was subsequently found to support electron uptake in the archaeon *Methanosarcina barkeri* ^10^, hinting that the underlying chemistry could be phylogenetically widespread. Critically, the demonstration in *S. epidermidis* relied on synthetic, pre-folded G4 oligonucleotides added exogenously. Whether bacteria assemble electroactive G4 structures themselves, and whether such natively produced structures are functional within an intact biofilm, has not been shown.

The question is timely because extracellular DNA and RNA are now recognised as structural constituents of the biofilm matrix, and both can adopt non-canonical conformations including G4; that stiffen the matrix and increase biofilm resilience^11–13^. G4 have been detected in pathogenic biofilms from clinical and model infections^11,14^ yet their physiological role beyond mechanical reinforcement is largely unexplored, as are the conditions under which biofilms form. If natively assembled matrix G4 are electroactive, they would couple the structural and metabolic states of a biofilm through a single class of molecule, which has profound implications for our understanding of how pathogens sustain themselves in the oxygen-limited interior of an infection.

We addressed this research gap in *Staphylococcus aureus*, a leading cause of bacteraemia, endocarditis, osteomyelitis and device-associated infections, and among the bacterial pathogens responsible for the greatest global mortality^15,16^. Its propensity to form recalcitrant biofilms at nutrient- and oxygen-limited infection sites makes it both a clinically important and a mechanistically informative model for asking whether natural G4-mediated EET occurs.

Here we show that *S. aureus* biofilms assemble G4-rich extracellular nucleic acids that bind hemin to form a catalytically active, electron-conducting complex without exogenous G4 addition. Accumulation of G4 correlates with biofilm age under nutrient-starvation rather than cell density. As biofilms develop, the G4 reorganise between reproducible spatial architectures, from intercellular networks to only being associated with the cell envelope. Combining electrochemistry, peroxidase imaging and structure-selective nuclease treatments, we find that these architectures impose distinct modes of electron transfer: diffusion-limited transfer through the matrix network versus surface-confined transfer at the cell envelope. Degrading the G4 abolishes electroactivity, whereas removing canonical B-DNA does not, establishing that the G4 fraction specifically carries the biofilm’s EET. Together these findings show that a pathogen builds and spatially organises its own electroactive nucleic-acid network. It also identifies biofilm architecture as a tunable determinant of extracellular electron flow and G4/hemin EET as a candidate target for destabilising the metabolism of recalcitrant infectious biofilms.

## Results

### Starvation induces natural extracellular G4 formation

We investigated natural G4 production in *S. aureus* biofilms grown under nutrient-limited conditions that could trigger mechanisms for release of extracellular DNA due to starvation stress in a dense bacterial population. After 4 d of growth without medium replenishment, biofilms exhibited abundant extracellular G4 structures detected by immunofluorescence with the G4-specific antibody BG4 (Fig. 1a). In contrast, biofilms with daily medium changes showed minimal G4 formation despite increased biomass and cell density (Figure 1b,c).

**Fig. 1.**
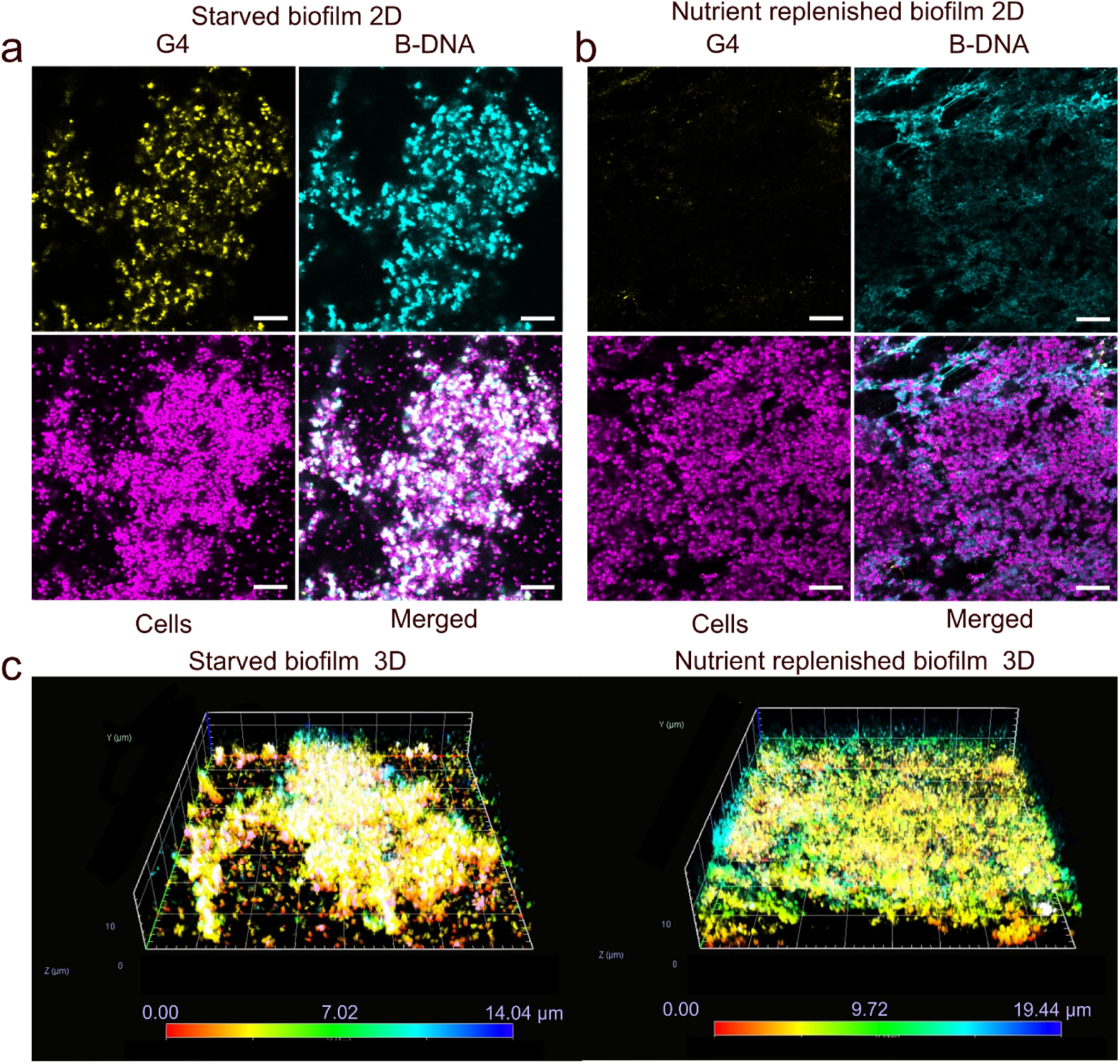
Starvation induces natural extracellular G4 formation in *S. aureus* biofilms. **(a)** A representative CLSM image of a 4-day old culture of *S. aureus* biofilm grown in BHI/NaCl with agitation at 150 rpm in a 200 µl culture volume without medium change. **(b)** A representative CLSM image of similar biofilm grown with daily replenishment medium replenishment. Biofilms were visualised as cells (magenta, FM 4-64 stain), immunolabelled G4s (yellow) and immunolabelled B-DNA (cyan). Scale bars = 10 µm **(c)** Colour gradient three-dimensional images indicate similar thickness but higher surface coverage for media replenished biofilms.

The stark difference between nutrient-starved and nutrient-replenished biofilms despite similar biofilm thickness indicates that G4 formation is triggered by metabolic stress rather than simply by biofilm age or cell density. This finding suggests that nutrient depletion, characteristic of chronic infection sites, may be a key environmental cue driving G4 assembly in pathogenic biofilms.

### Age-dependent restructuring of G4-DNA creates distinct biofilm developmental states

Having established that starvation induces G4 formation, we investigated whether biofilm age influences G4 spatial organisation. We extended incubation times from 3 to 12 d under continuous nutrient limitation and analysed G4 distribution patterns by immunolabeling and confocal microscopy. Remarkably, biofilms at different developmental stages exhibited reproducibly distinct G4 architectures that fell into three categories (Fig. 2).

**Fig. 2.**
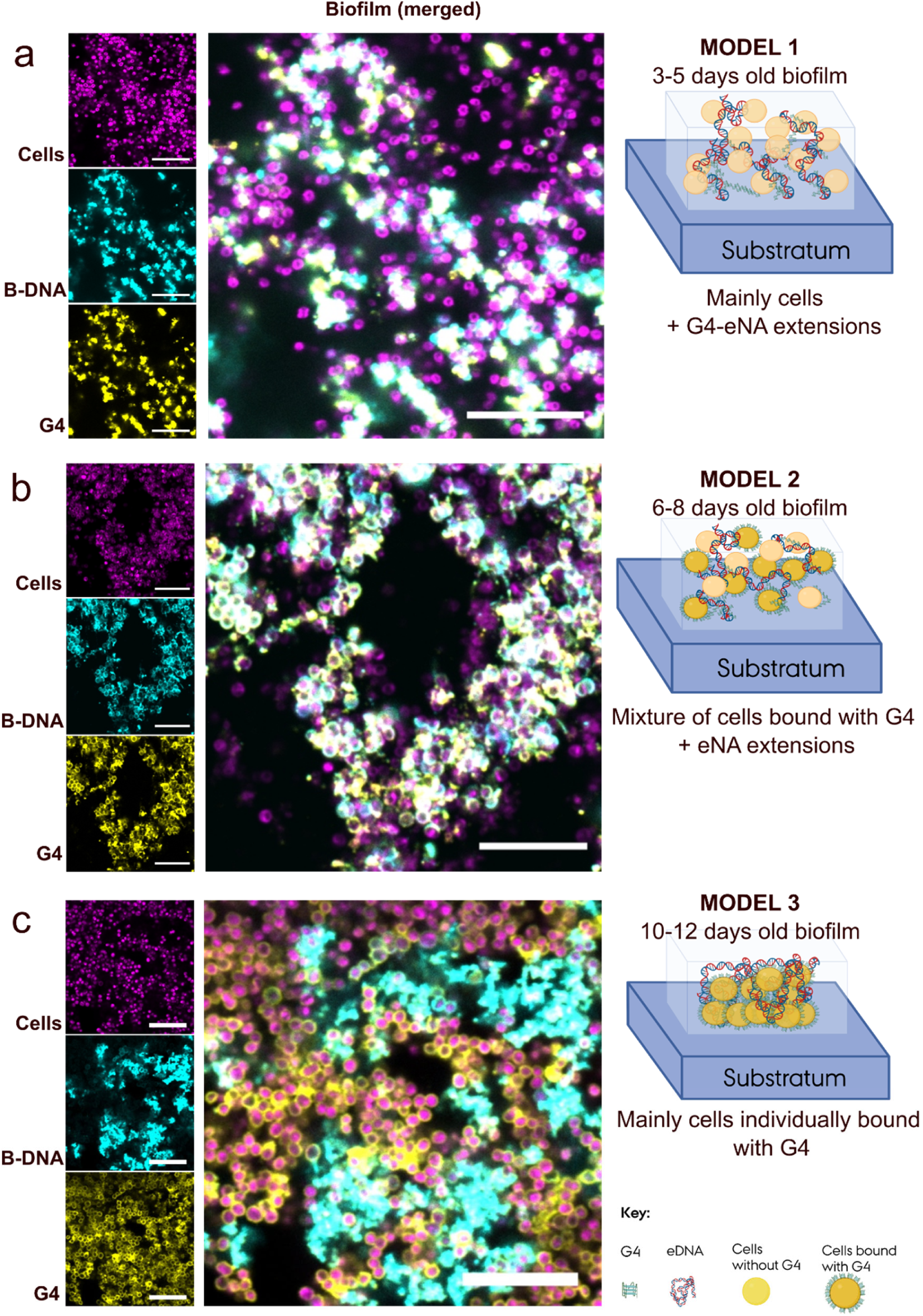
Arrangement of extracellular nucleic acids (eNA) progresses through a reproducible pattern as biofilms age. Representative biofilms grown for 3 - 12 d in BHI/NaCl without medium replenishment showed a reproducible pattern in distribution of extracellular G4, leading us to propose 3 distinct biofilm models. Model 1 **(a)** is 3-5 d old biofilm, Model 2 **(b)** is 6-8 d old biofilm, and Model 3 **(c)** is 10-12 d old biofilm. Biofilms were visualised as cells (magenta, FM 4-64 stain), immunolabelled G4s (yellow) and immunolabelled B-DNA (cyan). Scale bar = 10 µm. Schematic representation of orientation of cells and eNA arrangements show interactions between cells and G4 in the biofilm matrix. Schematic diagram created in BioRender. Ajunwa, O. (2026) https://BioRender.com/k8bt76m.

Young biofilms (3-5 d) displayed G4s as extracellular networks bridging clusters of cells, with G4 structures at the bacterial envelope, but also intertwined with B-DNA and extending beyond cell surfaces (Model 1, Figure 2a). These network configurations suggest that newly released genomic DNA initially organises into extended scaffolds connecting cell aggregates. Intermediate-age biofilms (6-8 d) showed a transitional, heterogeneous architecture. Some cells retained network-associated G4s while others exhibited G4 associated with the cell envelope (Model 2, Figure 2b). This mixed state implies active reorganisation of extracellular nucleic acids during biofilm maturation.

Mature biofilms (10-12 d) underwent an architectural transformation with G4 predominantly associated with the cell envelope, while B-DNA remained spatially separated from these (Model 3, Figure 2C). If G4 are G4-DNA integrated in a DNA superstructure, we would have expected G4 and B-DNA to co-localise. The physical separation might suggest that G4 is G4-RNA, and we test this directly below (Fig. 7, Extended Data Fig. 1).

This developmental progression from intercellular networks to envelope-associated G4 indicates that biofilm maturation involves remodelling of the extracellular nucleic acid architecture. The transition is consistent with an active process rather than passive accumulation: extracellular DNA is initially a connecting scaffold, but then becomes redistributed to the cell envelope. Alternatively, the envelope-associated G4 are secreted at this stage in biofilm formation through processes that are distinct from other mechanisms for producing extracellular DNA and RNA.

### Natural G4s in *S. aureus* biofilms form a catalytically active complex with hemin

G4 can bind hemin *in vitro* and form a DNAzyme with peroxidase-like activity^8^, but the catalytic activity depends on the G4 structure and neighboring nucleotide sequences^17^. We previously showed that catalytically active G4/hemin can be used as a conduit for extracellular electron transfer in bacteria^1^. If naturally formed G4 provide catalytic activity when interacting with hemin, they have potential to fundamentally impact the biology of *S. aureus*.

Using tyramide signal amplification to detect the location of extracellular peroxidase activity, we confirmed that all three biofilm architectures are catalytically active when supplemented with hemin (Fig. 3). Strikingly, despite their distinct spatial organisations, all models exhibited similar catalytic activity concentrated at the cell envelope, indicating that catalytically active G4 are primarily located here.

**Fig. 3.**
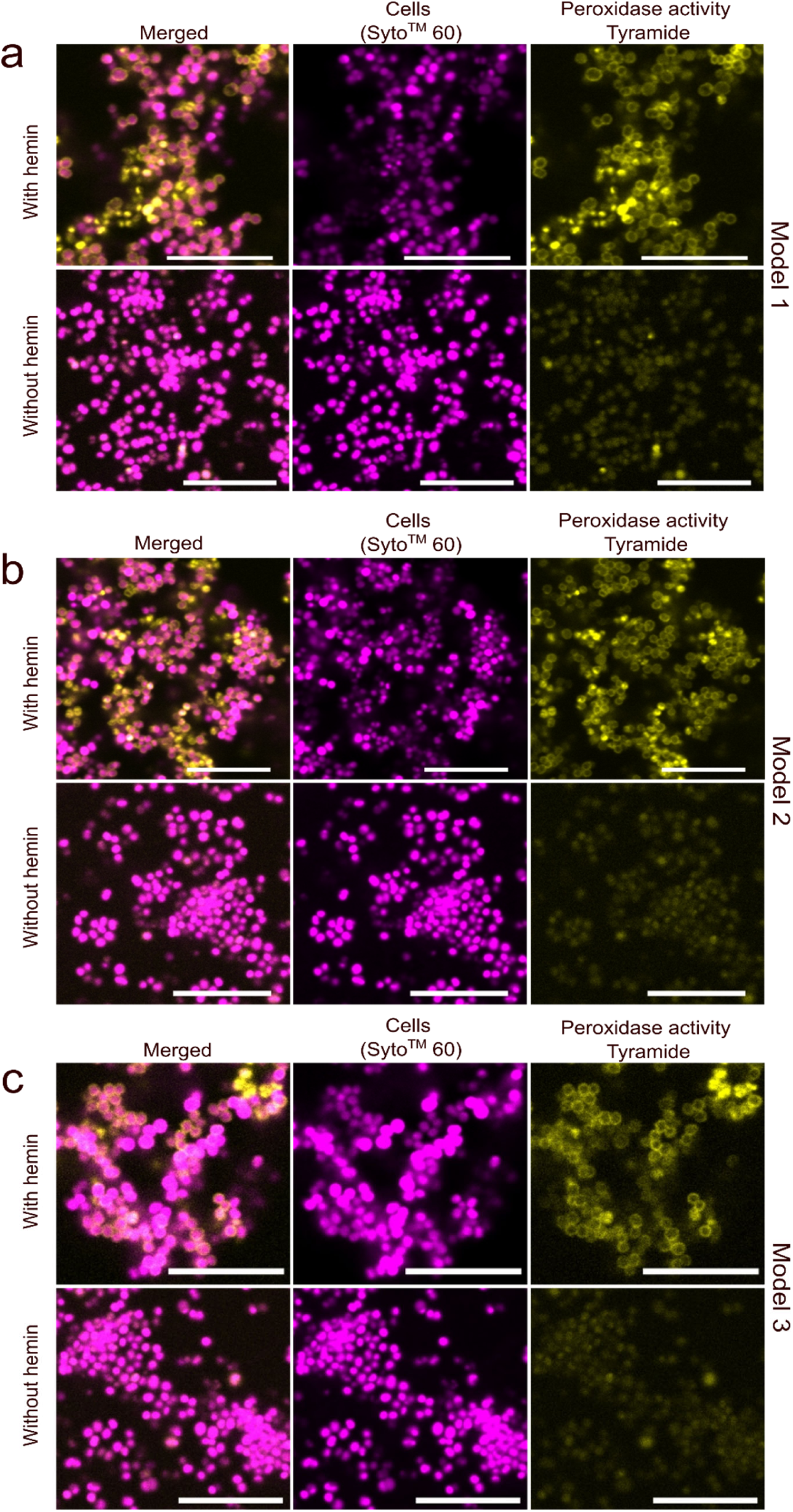
Naturally occurring G4 display peroxidase activity in the presence of hemin. Biofilms were visualised by staining cells (magenta, SYTO60^TM^) and extracellular peroxidase activity (yellow, Tyramide Signal Amplification). Scale bar = 10 µm. **(a)** Model 1, **(b)** Model 2, and **(c)** Model 3. Images are representatively chosen from 3 replicates.

### Natural G4s interact with hemin resulting in varying EET mechanisms

Each biofilm model generated a characteristic electrochemical signature as shown by cyclic voltammetry (CV) and differential pulse voltammetry (DPV). The three biofilm models were first grown under aerobic conditions on screen-printed electrodes without any electrode potential. The spent medium was then replaced with fresh medium (supplemented with 10 µM hemin) and incubated under anoxic conditions with electrode potential (poised at +400 mV), establishing conditions under which electron transfer to the electrode could sustain anaerobic metabolism. DPV analysis was conducted after 24, 48 and 72 h to allow time for the biofilm to establish EET-mediated metabolic activity via the electrode.

The biofilms became electroactive, and CV analysis showed oxidation and reduction curves with significantly higher currents than those acquired in sterile medium (Extended Data Fig. 2). The gradual increase in electroactivity was reflected in increasing DPV peaks during the 72-h incubation on the poised electrode (Fig. 4), and subsequently the current production remained stable for at least additional 50 h (Extended Data Fig. 3). Model 1 biofilms (network-associated G4s) exhibited a dominant reduction peak at approximately -0.18 V, consistent with G4-hemin complexes within extracellular DNA networks mediating electron transfer^1^. Model 3 biofilms (cell-envelope G4s) showed a prominent oxidation peak at approximately +0.28 V, characteristic of membrane-associated redox species. Model 2 biofilms displayed both signatures as a convoluted peak ranging from -0.15 V to +0.2 V during the 24 to 72 h rise in metabolic activity, confirming its hybrid architecture. These distinct electrochemical profiles indicate that the spatial organisation of G4 structures is associated with distinct modes of EET.

**Fig. 4.**
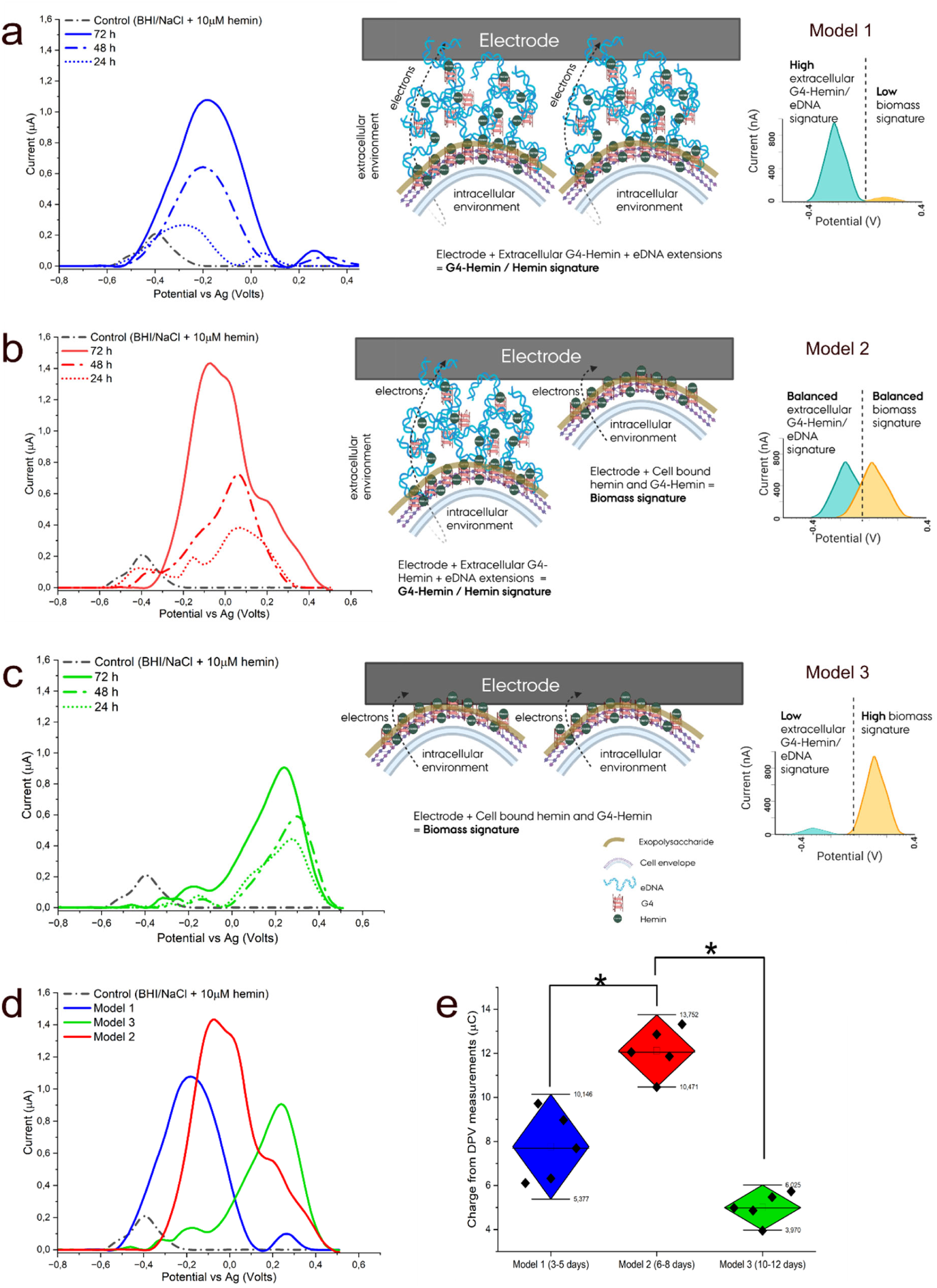
Electrochemical profiling of biofilms with different G4 arrangements revealed distinct electrochemical signatures for matrix-associated vs envelope-associated G4/hemin. Electrochemical signals showed a gradual metabolic adaptation of the starved biofilms to EET as electron flow increased over 72 h in all biofilm models. DPV curves are averages of n > 3 biological replicates. Curves were normalised on PSTrace® software to remove signals from non-Faradaic currents, leaving only signals from Faradaic currents. Sterile BHI/NaCl medium supplemented with 10 µM hemin was used as control. **(a)** Model 1, **(b)** Model 2, **(c)** Model 3, **(d)** A combined plot of electrochemical potentials of all three models. **(e)** Charge values were obtained by integrating current from DPV measurements based on the area under each curve for each biofilm model. One way ANOVA and Tukey’s test - asterisks above the bars indicate significant differences (\**p* < 0.05). Schematic diagrams were created in BioRender. Ajunwa, O. (2026) https://BioRender.com/v4h0j80

Control samples with sterile media supplemented with hemin had an electrochemical peak at -0.38 V resulting from dissolved hemin (Fig. 4, Extended Data Fig. 2a) while hemin-deficient media or media supplemented with the iron-deficient protoporphyrin displayed no peak (Extended Data Fig. 2). These controls validate that hemin is needed for *S. aureus* electroactivity, and that the electrochemical profiles of *S. aureus* biofilm can be ascribed to immobilised hemin and not just from dissolved hemin.

To confirm that biofilm architectures remained stable during electrochemical measurements, we imaged biofilms directly on electrodes after 72 h of anaerobic incubation (Figure 5). G4 arrangements were preserved in all three models, confirming that the distinct electrochemical signatures reflect structural differences rather than electrode-induced reorganisation.

**Fig. 5.**
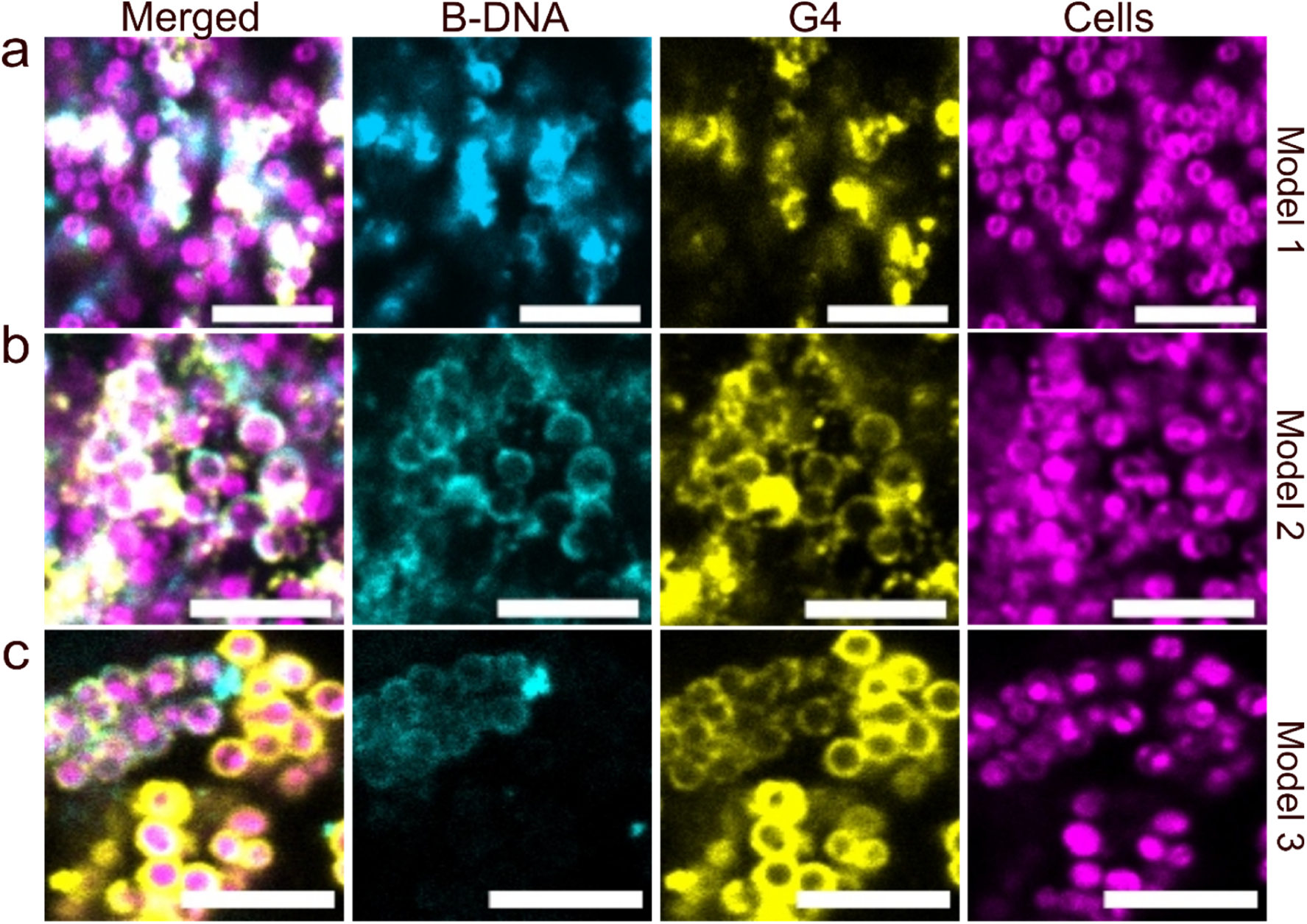
G4-rich biofilm models retain cell-G4-eNA arrangements under electroactive conditions. Microscopy of electrode surface after electrochemical measurements validate that the same structural arrangements of biofilms formed from starvation at different incubation times remained intact during and after EET measurements. Scale bar = 5 µm **(a)** Model 1, **(b)** Model 2 and **(c)** Model 3.

### G4 proximity to cell membranes tunes electron transfer mode

The distinct electrochemical signatures of the three biofilm models suggested fundamentally different mechanisms of electron transfer. To understand how G4’s spatial organisation relates to EET, we examined the relationship between G4-cell distances and electron transfer kinetics.

Confocal microscopy revealed systematic differences in G4 localisation across models (Figure 6A, B). In Model 1, G4 structures formed networks extending from cell surfaces, intertwined with extracellular B-DNA. Model 3 showed the opposite arrangement: G4s associated tightly to the envelope of individual cells (0.3-0.6 μm from membrane), with minimal network extensions. Model 2 exhibited both configurations within the same biofilm, consistent with its intermediate developmental state.

**Fig. 6.**
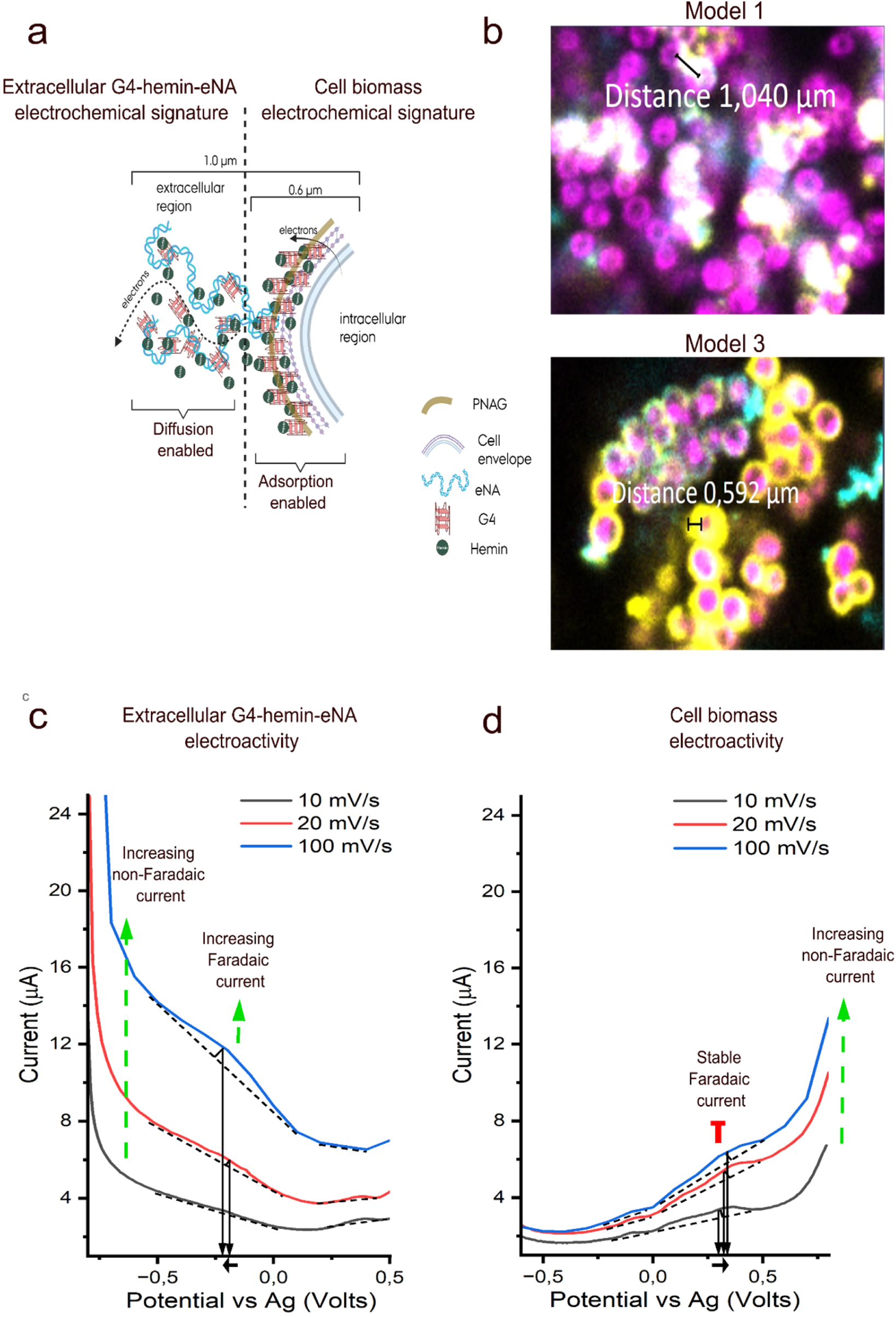
Spatial organisation of G4-NA determines electron transfer mechanism. (A) Schematic illustration of two distinct architectures: network-associated G4 (Model 1) and cell envelope-associated G4 (Model 3). (B) Representative CLSM images showing differences in the location of G4 in Model 1 and 3 biofilms. (C) Current-Potential plots of Model 1 biofilms at scan rates of 10, 20, and 100 mV/s. Faradaic current (peak at - 0.18 V) increases with scan rate, hinting at a diffusion-enabled electron transfer through extracellular DNA networks. (D) Current-Potential plots of Model 3 biofilms at scan rates of 10, 20, and 100 mV/s. Faradaic current (peak at +0.28 V) remains stable across scan rates, indicating a surface-confined electron transfer via direct electrode contact with G4-bound cells. With increasing scan rates, there was a shift in peak midpoints in curves to the negative potential for extracellular G4-hemin-eNA electroactivity and to the positive potential for cell biomass electroactivity. In both curves the data was analysed on PSTrace and was an average of 3 biological replicates. Curves are not baseline-corrected to show the effects on capacitive (non-Faradaic) currents. Schematic diagram was created in BioRender. Ajunwa, O. (2026) https://BioRender.com/b5c67sj

To determine whether these architectural differences affect electron transfer, we performed voltammetry at increasing scan rates on Models 1 and 3. The scan rate experiments revealed distinct behaviours from the two architectures (Figure 6C, D). Model 1 showed scan-rate-dependent increases in faradaic current, indicating diffusion-dependent electron transfer, particularly with negative potentials (approx. -0.18 V). This phenomenon occurs when redox-active species are distributed throughout a porous matrix and diffuse to the electrode surface to transfer electrons. In contrast, Model 3 faradaic currents were stable and independent of the scan rate at positive potentials (approx. +0.28 V), characteristic of surface-confined electron transfer. Here, we propose that immobilised G4/hemin interacts with cell surface proteins and facilitate direct electrode contact and transfer electrons without requiring diffusion. The interaction could explain the different electrochemical signature from the negative potential observed in model 1.

### Enzymatic degradation implies electron transfer via G4-RNA

The correlation between G4 spatial organisation and electrochemical signatures suggested that G4 structures directly mediate electron transfer. However, biofilm matrices contain diverse nucleic acid species: canonical B-DNA, non-canonical structures, and RNA. To determine whether G4s specifically enable EET, or whether other extracellular nucleic acids contribute, we enzymatically dissected the biofilm matrix using nucleases with different structural specificities. We treated electroactive biofilms with four enzymes: DNase I (cleaves dsDNA (B-DNA)), micrococcal nuclease (cleaves dsDNA and dsRNA, including G4 structures), S1 nuclease (cleaves ssDNA and ssRNA), and RNase A (cleaves ssRNA). After 72 h of electroactive growth, biofilms were exposed to each enzyme, and electrochemical activity was reassessed by differential pulse voltammetry (Fig. 7).

**Fig. 7.**
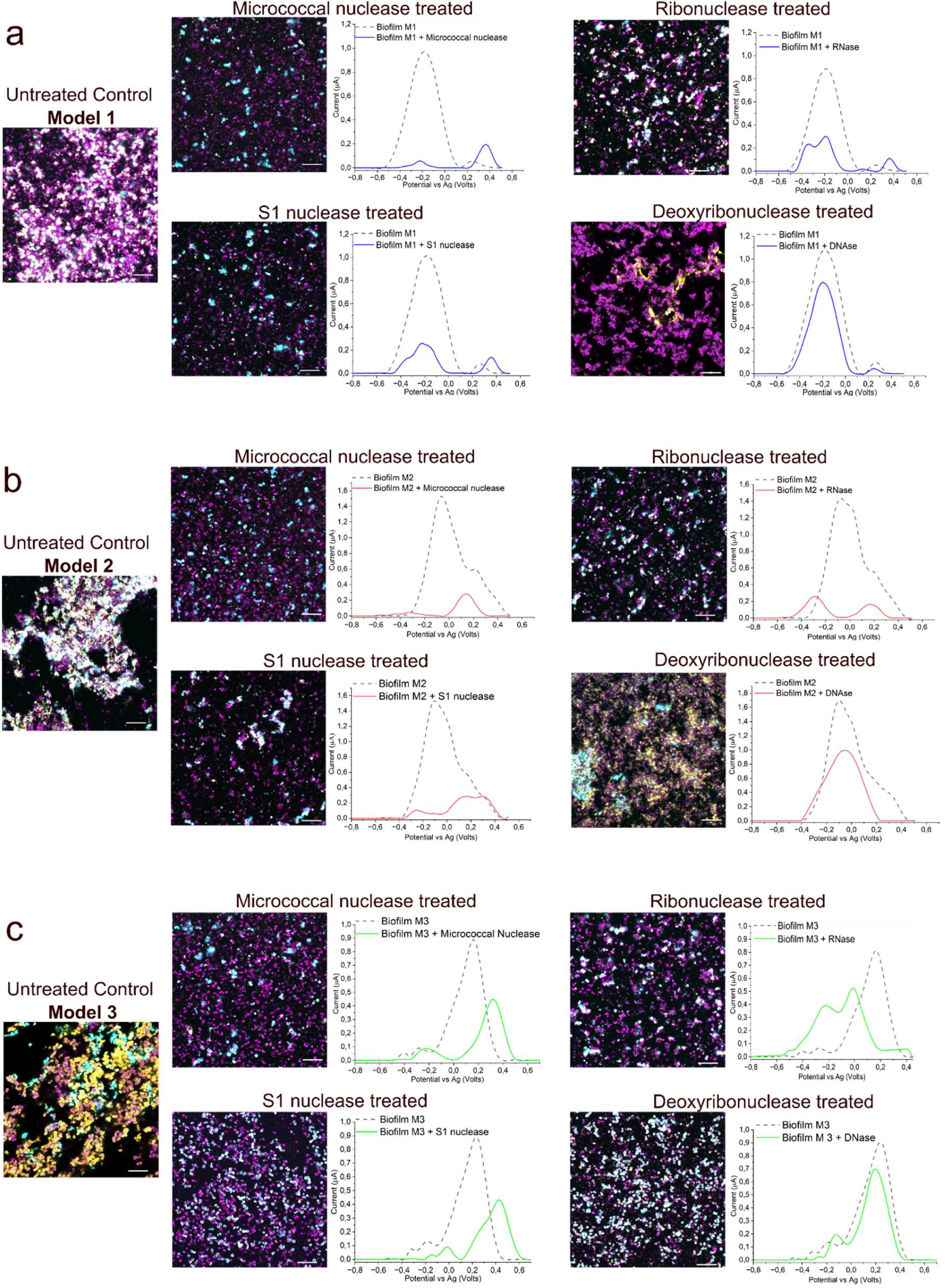
Enzymatic cleavage reveals G-quadruplex structures are essential for electroactivity and biofilm cohesion. Electroactive biofilms (Models 1-3) were treated with nucleases targeting different nucleic acid structures, then reassessed by differential pulse voltammetry and confocal microscopy. Scale bars = 10 μm. **(a)** Model 1 (network architecture). DNase I treatment minimally affected electrochemical signature. Microscopy shows enzymatic treatment dispersed cell clusters and reduced electrode attachment. **(b)** Model 2 (hybrid architecture). Nuclease treatment partially deconvoluted the broad electrochemical peak, revealing underlying components. Cell-DNA network connections were visibly disrupted. **(c)** Model 3 (envelope-associated architecture). RNase A uniquely altered the electrochemical signature, shifting the dominant peak from positive (+0.28 V) to negative potentials, consistent with removal of G4 from cell envelopes. DPV curves show representative biological replicates.

DNase I treatment caused minimal reduction in electron transfer across all three models, with peak position being unchanged and exhibiting minimal reduction in DPV current (Fig. 7). Converting DPV currents to charge, DNase I caused only a 10-22% charge reduction (Figure 8). In contrast, micrococcal nuclease dramatically reduced electroactivity in all models with up to 88% charge reduction (Fig. 8), and both S1 nuclease and RNase A resulted in similar effects. Collectively, these results indicate that the nucleic acid structures involved in electroactivity include RNA and non-canonical structures that are resistant to DNase I, such as G4. Interestingly, RNase A treatment not only reduced the current, but also shifted the dominant peak from positive potentials (+0.28 V, characteristic of cell-surface electron transfer) to two peaks, one at 0 V and a negative potential at -0.25 V, characteristic of potentials within the range of network-mediated transfer (Fig. 7c). This shift also suggests that RNA structures contribute to cell-surface electroactivity in mature biofilms.

**Fig. 8.**
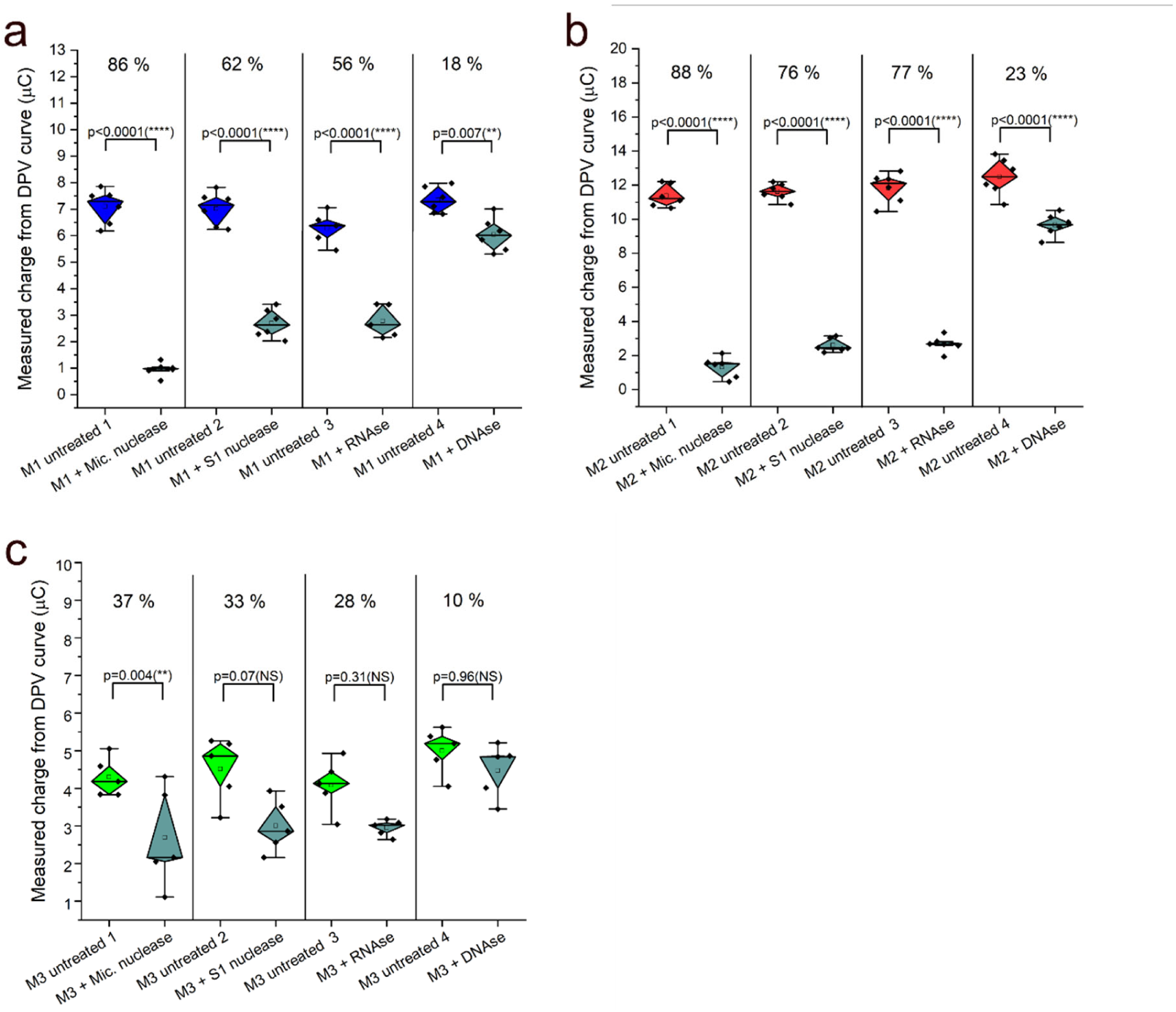
Percentage reduction in average electrical charge from DPV measurements for models 1-3 (M1-3). **(a)** Enzyme treatment of Model 1 biofilms revealed a moderate impact on electroactivity by DNase I (18% charge reduction) and a large impact of micrococcal nuclease, and some impact of S1 nuclease, and RNase A (up to 86% charge reduction). **(b)** Treatment of Model 2 biofilms also confirmed that micrococcal nuclease had the largest impact on electroactivity (88.38 % charge reduction). **(c)** Model 3 biofilms were less affected by enzyme treatments with only a significant effect from micrococcal nuclease (37 % charge reduction). Charge reduction percentages represent mean ± SE from n>3 biological replicates. One way ANOVA and Tukey’s test - asterisks above the bars indicate significant differences (\**p* < 0.05, \*\**p* < 0.01, *** *p* < 0.001, *** *p* < 0.0001) while *ns* indicates lack of statistically significant difference

Beyond reducing electrochemical signals, all nuclease treatments disrupted biofilms, resulting in fewer bacteria on the electrodes (Fig. 7). The biofilm-disruptive effect of DNase I with minimal impact on electroactivity emphasises that nucleic acids impact both biofilm structure and electroactivity – but different structures can play different roles.

## Discussion

The discovery that synthetic G4-DNA complexed with hemin can act as an extracellular electron transfer (EET) conduit established a proof of concept^1^,but left open whether bacteria exploit this chemistry using nucleic acids they produce themselves. Here we show that *S. aureus* biofilms assemble such structures naturally, without exogenous G4 addition, and that the resulting electroactivity is organised in space by the biofilm’s developmental state. These findings connect three previously separate lines of work; eDNA-supported, mediator-based EET in *Pseudomonas aeruginosa*^4^, the structural and DNAzyme properties of biofilm G4^11^, and G4/hemin-mediated EET with synthetic oligonucleotides^1^ into a single phenotype that emerges from native matrix assembly. To our knowledge, this is the first demonstration that the spatial arrangement of cells and G4-rich extracellular nucleic acids (eNAs), rather than the mere presence of G4, determines how electrons move through a pathogenic biofilm.

Extracellular G4 accumulation was triggered by nutrient starvation. Starved biofilms were G4-rich, whereas age-matched biofilms with daily medium replacement remained G4-poor despite equal or greater biomass (Fig. 1). The extracellular localisation of these structures points to cell lysis as a likely source^18^, with released genomic DNA refolding into G4 once guanine-rich tracts encounter stabilising cations^11^, consistent with the salt- and shear-dependence of biofilm G4 formation reported previously ^11^. Spatially patterned lysis during starvation is well documented in other biofilm formers, where ordered lysis deposits eDNA and reshapes the matrix^19^ and released intracellular contents can act as signals that further modulate community structure^20^.

Across 3–12 d of starvation, G4 organisation fell into three reproducible architectures; intercellular networks (young), a mixed state (intermediate), and cell envelope-associated (mature) (Fig. 2). Because these were captured in separate biofilms rather than by time-lapse imaging, we interpret them as developmental states rather than a directly observed trajectory, and whether a single biofilm progresses through all three remains to be tested. The redistribution from extended networks to envelope-bound is nonetheless consistent with active matrix remodelling, plausibly by secreted nucleases. Extracellular nucleic acids can be dynamically turned over in biofilms^21^, and micrococcal nuclease, a native *S. aureus* enzyme, has been shown to be a very potent disruptor of both architecture and electroactivity in our experiments (Fig. 7, 8). The role of self-secreted nucleases in the network-to-envelope transition remains to be established.

The central functional result is that each architecture imposed a distinct EET mechanism, attributable to the position of catalytically active G4/hemin relative to the cell and electrode. Hemin was the limiting redox component: the iron-free analogue protoporphyrin IX supported no electroactivity (Extended Data Fig. 2), confirming that the signals arise from G4/hemin rather than from G4 or porphyrin alone. In network-dominated young biofilms (Model 1), G4/hemin distributed through eDNA produced scan-rate-dependent faradaic currents at negative potential the diagnostic signature of a diffusion-limited process in which mobile redox species must reach the electrode. In mature biofilms (Model 3), envelope-associated G4/hemin produced scan-rate-independent currents at positive potential, characteristic of surface-confined transfer through immobilised complexes in directly in contact with the electrode, while the intermediate architecture (Model 2) displayed both. These regimes represent a trade-off: diffusion-based transfer is fast but depends on cell-to-electrode distance and is vulnerable in thick biofilms or upon detachment, whereas surface-confined transfer is distance-independent and stable but limited by the electrode-contacting cell area ^22^. The positive potential of the surface-confined regime agrees with the redox range of cell-envelope components, including membrane cytochromes^23^, and with early voltammetry of electrode-attached biofilms^24^, suggesting that envelope-bound G4/hemin may couple to membrane redox partners. That EET mode tracks the matrix gradients and structural heterogeneity intrinsic to biofilms^25^ indicates that architecture is a tunable determinant of electron flow rather than an incidental correlation.

Cell envelope-localised peroxidase activity in all three models (Fig. 3) correlated with architecture-specific electrochemistry. Catalytically active G4/hemin at the cell envelope is common to every model, validating surface peroxidase signals regardless of EET mode. Tyramide deposition reflects local peroxidase activity at a fixed point, whereas voltammetry integrates electron flux across the entire biofilm–electrode interface and is therefore additionally sensitive to the network-distributed G4/hemin that underlies the diffusion-limited signal in young biofilms. Imaging and electrochemistry confirmed catalytically active envelope-associated G4/hemin in every model, and resolving the additional network-associated fraction that distinguished young biofilms.

Beyond defining the electron transfer mode, native G4/hemin allowed starved biofilms to switch from aerobic to anaerobic, electrode-dependent metabolism when supplied with fresh nutrients and hemin (Extended Data Fig. 4, 5). This supports the hypothesis that metal-porphyrin/G4 chemistry is a generic route for microbial survival under oxygen limitation, now demonstrated for natively produced nucleic acids. The dependence on nucleic acids is notable because the same eNAs that scaffold the matrix and serve as a reclaimable phosphate and nutrient reservoir^21^ also carry the biofilm’s electroactivity, coupling structure, nutrition and electron flow through a single class of molecule. Our enzyme experiments make this coupling explicit: DNase I removed canonical B-DNA and dispersed cells with little effect on current, whereas G4- and RNA-targeting treatments abolished electroactivity (Figs. 7, 8), dissociating the structural role of bulk eDNA from the electroactive role of G4 and associated RNA.

The conditions that triggered G4 assembly: nutrient depletion, high cell density and oxygen limitation, are characteristic of the chronic and device-associated infections in which *S. aureus* is most difficult to treat. We previously detected G4 in staphylococcal biofilms recovered from a murine implant-associated osteomyelitis model^11^, and G4 have been reported in airway infections^26^ and in oral biofilms, where they associate with bacterial surfaces as observed here^14^. In an infection, the hemin supplied here exogenously would instead be derived from host heme scavenged by *S. aureus*, and the poised electrode that served as the terminal acceptor would be replaced by physiological acceptors - oxygen at the aerobic biofilm interface, host redox-active surfaces, or hydrogen peroxide via the peroxidase activity of the G4/hemin DNAzyme^1,11^. G4-mediated EET may therefore help sustain low-level metabolism in the anoxic interior of an infectious biofilm.

Taken together, *S. aureus* biofilms naturally assemble G4-rich eNAs that bind hemin to form a redox-active DNAzyme, and the spatial arrangement of these complexes set by starvation and developmental state, determines whether electrons move by diffusion through the matrix or by surface-confined transfer at the cell envelope. Owing to the fact that this chemistry requires only guanine-rich nucleic acids and an abundant metal-porphyrin^1,10^, it may represent a broadly conserved strategy for energy conservation under oxygen limitation. The dependence of biofilm electroactivity on G4 integrity, and its sensitivity to a native staphylococcal nuclease, identifies G4/hemin EET as a candidate target for strategies aimed at destabilising the metabolism of recalcitrant infectious biofilms.

## Methods

### Bacterial strains and culture conditions

*Staphylococcus aureus* ATCC® 29213™ was utilised throughout this study, and the strain was stored at -80°C as a 30% glycerol stock tube culture before inoculation. Brain heart infusion broth (BHI, Merck) and BHI agar were used in this study unless otherwise stated. BHI was supplemented with 0.2 M NaCl for preparing overnight cultures, inoculated from single colonies into 20 ml broth and incubated for 15–20 h at 37°C with 150 rpm shaking.

### Preparation of biofilms

*S. aureus* was harvested from overnight cultures by centrifugation (5 min at 906 × g) and resuspended in BHI/NaCl to an optical density (OD_600_) of 0.01 before transferring 200 µl to 96-well plates (Ibidi Treat µ-Plate^TM^) and incubating in a humid atmosphere (ziplock bag) at 37°C and 150 rpm shaking for 3 – 12 d without replenishing the media. Three biofilm models were established with different durations of the incubation - Model 1: 3–5 d, Model 2: 6–8 d, and Model 3: 10–12 d. For comparison, biofilms were also grown for 3 d with daily medium change.

### Immunolabelling and biofilm visualisation

Structure-specific antibodies were used to visualise extracellular nucleic acids in the biofilms. We visualised G4 with Atto488-conjugated BG4 antibody (1 mg/ml stock in phosphate-buffered saline, PBS) (goat monoclonal IgG lambda, Ab00174-24.1, Absolute Antibodies®). The B-DNA-specific antibody AB1 (1mg/ml stock in PBS) (mouse monoclonal, AB27156, Abcam®) was not conjugated and therefore required labeling with Atto 405-conjugated secondary anti-mouse antibody AB2 (2 mg/ml stock in PBS with 50% glycerol and 0.05 % sodium azide) (goat monoclonal IgG H + L, A-31553, Invitrogen®).

Immunolabeling was performed by diluting antibodies, 1:100 in 3 % bovine serum albumin (BSA) in PBS and incubating them with the sample as follows: Biofilms in 96-well plates were gently washed in sterile distilled water before adding 60 µl antibody solution and incubating for 2 h with mild shaking (40 rpm) at room temperature (RT). When multiple antibodies were used together, the second antibody (e.g. AB1 following BG4) was added by subsequently adding 60 µl of antibody solution and incubating the sample for additional 1.5 h. The solutions were then decanted, and biofilms were washed gently with 3% BSA in PBS. For AB1 labeling, we added a third hybridisation step for the secondary antibody by adding 60 μl of the secondary antibody AB2 to the washed biofilm and incubating for 60 min. Biofilms were finally washed in 100 mM NaCl.

Bacterial membranes were then labeled with FM 4-64 (T13320, Invitrogen), diluting at 1 mg/ml stock solutions 10 μg/ml working concentration in 100 mM NaCl, and incubating the samples for 15 min. The solution was then aspirated, and biofilms were mounted with Prolong^TM^ Antifade solution (Invitrogen, P3698). Imaging was performed with a confocal laser scanning microscope (CLSM) (LSM900, Carl Zeiss) equipped with a 63× objective (Plan-Apochromat, NA1.4). Acquisition settings were 488 nm excitation and >660 nm emission to visualise bacteria stained with FM 4-64, while 405 excitation and 410– 477 nm emission for the Alexa Fluor 405-conjugated AB2, and 488 nm excitation and 520–600 nm emission for Atto488 conjugated BG4. Image analysis of 2D and 3D images were done on the ZEN® 3.7 (ZEN lite) software.

### Visualisation of G4-hemin oxidoreductase activity

Biofilms were prepared as stated above and gently washed in 200 μl modified MES buffer [25 mM MES (Sigma, M3671), 0.2 M NaCl (Sigma, S5886), and 0.010 M KCl (Sigma, 1.04936)] adjusted to pH 6.5. Biofilm samples were then supplemented with and without 10 μM hemin (from 10 mM stock in 200 mM Tris, 30% DMSO, and 100 mM NaOH, pH 11, stored in the dark) and incubated at room temperature for 30 min with mild shaking (50 rpm). The buffer was then gently aspirated, and biofilms were incubated for 90 min (50 rpm, RT) with 90 μl tyramide reagent containing Alexa Fluor 488-conjugated Tyramide (1:100 dilution) (Invitrogen, B40953), 2 mM ATP (Thermo Scientific, R0441), and 0.1% hydrogen peroxide (Sigma, 216763) in modified MES buffer. After incubation, the solution was aspirated and washed with modified MES buffer before staining the biofilm with 90 μl 20 μM SYTO^TM^ 60 (Invitrogen, S11342) in deionised water. The solution was aspirated, and antifade solution (Invitrogen, P3698) was added before CLSM imaging. Alexa Fluor-488-conjugated tyramide was visualised by 488 nm excitation and <550 nm emission. SYTO^TM^ 60-stained bacteria were visualised by 647 nm excitation and 650 – 700 nm emission.

### Electroanalyses of biofilm models

Overnight cultures of *S. aureus* were harvested by centrifugation, resuspended in BHI/NaCl to an OD_600_ of 0.05, and 0.4 ml was transferred to custom-made 1 ml glass tubes mounted on screen printed electrodes (SPEs, Metrohm DropSens DRP-ITO10-U20, Metrohm Spain) using non-conductive glue, adapted from the set-up by^27^. Tubes had butyl rubber caps for easy syringe piercing. SPEs had a 4 mm diameter transparent Indium Tin Oxide (ITO) working electrode (WE) with surface area of 0.126 cm^2^, a graphite counter electrode, and Ag pseudo-reference electrode. Prior to use, tubes with SPEs were sterilised in 70% v/v ethanol, washed thrice in sterile deionised water, and air dried. After inoculation, samples were incubated aerobically at 37°C for duration specified for each biofilm model. The liquid was then aspirated and replaced with BHI/NaCl supplemented with 10 µM hemin and gently sparged with N_2_ for 2 min to remove oxygen. Biofilms were then incubated statically at 37°C with electrodes poised at +0.4 V for up to 72 h. Control samples contained sterile medium with 10 µM hemin. Hemin-free biofilm controls were either incubated in BHI/NaCl without hemin or supplemented with 10 µM iron-deficient protoporphyrin (IX). To determine the influence of nutrient and hemin supplementation on metabolic recovery rates, further controls without any media replenishment were analysed. Electrochemical analyses were carried out during and after 72 h biofilm growth.

Electrodes were connected to a computer-controlled multichannel PalmSens 4 potentiostat (PalmSens® Netherlands). Electrochemical data were collected using the PalmSens PsTrace 5.9 software, and cyclic voltammetry (CV), differential pulse voltammetry (DPV), chronoamperometry (CA) and charge measurements were conducted. CV was run between potentials -0.8 and 0.8 V vs Ag at a scan rate of 0.05 V/s and step potential of 0.05 V. Three scan cycles were performed, while the last scan was selected. DPV analyses were performed between -0.8 V as start potential and +0.8 V stop potential with scan rate of 0.05 V/s. The average scans of three biological replicates were used for each experiment. For varied scan rate DPV analyses, three different scan rates were applied: 0.01 V/s, 0.02 V/s and 0.1 V/s. Voltages of scan rates and step potentials were kept at a constant 1:1 ratio for each scan rate. Time dependent CA analyses were carried out after electroactive growth of the biofilm for 72 h at poised potential of + 0.4 V. Same poised potentials were still used to capture all possible EET signals generated from the different biofilm models. For short term CA analyses, after 72 h growth, biofilms were further monitored for additional 20,000 s (5.6 h), while for long term CA analyses biofilms were monitored for 180,000 s (50 h). At specific time points during short term and long-term CA measurements, charge values in coulombs (C) were obtained by converting current as a function of time. Also charge data were recorded for DPV analyses to determine specific charge generated from DPV curves by integrating the areas under each DPV curve analysed. For electrode imaging, test electrodes were carefully detached from the base of the tube and immunolabeling as well as FM 4-64 staining were carried out on biofilm formed on the working electrode area of the ITO electrodes prior to CLSM image acquisition. Imaging was performed using same labeling and image acquisition settings detailed above.

### Nuclease treatments and electroanalytic re-testing

Nucleases were used in the following concentrations according to the method of Minero et al. ^11^: 500 U/ml DNase I, 15 U/ml micrococcal nuclease, and 25 U/ml S1 nuclease. RNAse A was used at 200 µg/ml, following modifications to the method of ^28^. Enzymes were prepared in buffer (25 mM Tris with 6.25 mM CaCl_2_ and 1 mM MgCl_2_, pH 6) shortly before application. 72 h old biofilms grown on SPE were treated with 200 µl enzyme solution at 37 °C for 2 h. An untreated control was included for every enzyme treatment. Treated and untreated biofilms were then analysed by repeating electrochemical and CLSM analyses following protocols stated above. Differences in DPV curves and charges recorded from area under DPV curves between treated and untreated samples were recorded. Enzyme solutions were aspirated and replaced with sterile BHI/NaCl with 10 µM hemin before analyses to create similar electrolyte conditions with untreated controls.

### Statistical and data analyses

Minimum of three independent biological repeats were carried out for each experiment unless otherwise stated. Data analyses and statistical analyses were done in Origin (Pro), version 2024. Electrochemical data were obtained and processed in PSTrace 5.9. Statistical analyses involved mean comparisons and one-way analyses of variance (ANOVA) based on Tukey’s post hoc tests and were applied for the determination of significant differences between pairs of means. Descriptive statistical analyses involved the use of box plots, means, medians and standard deviations in some graphs.

## Acknowledgements

We gratefully acknowledge the Independent Research Fund Denmark (grant no. 2032-00294B) and the Carlsberg Foundation (CF23-1540) for funding this work and give special thanks to Center for Electromicrobiology for sparring along the way.

## Author contributions

**OMA:** Conceptualisation, investigation, methodology, data curation, data analyses, writing – original draft, writing – review and editing

**RLM:** Conceptualisation, fund acquisition, methodology, writing – original draft, writing – review and editing

## Competing interest statement

Both authors declare no competing interests.

## EXTENDED DATA

**Extended Data Fig. 1:**
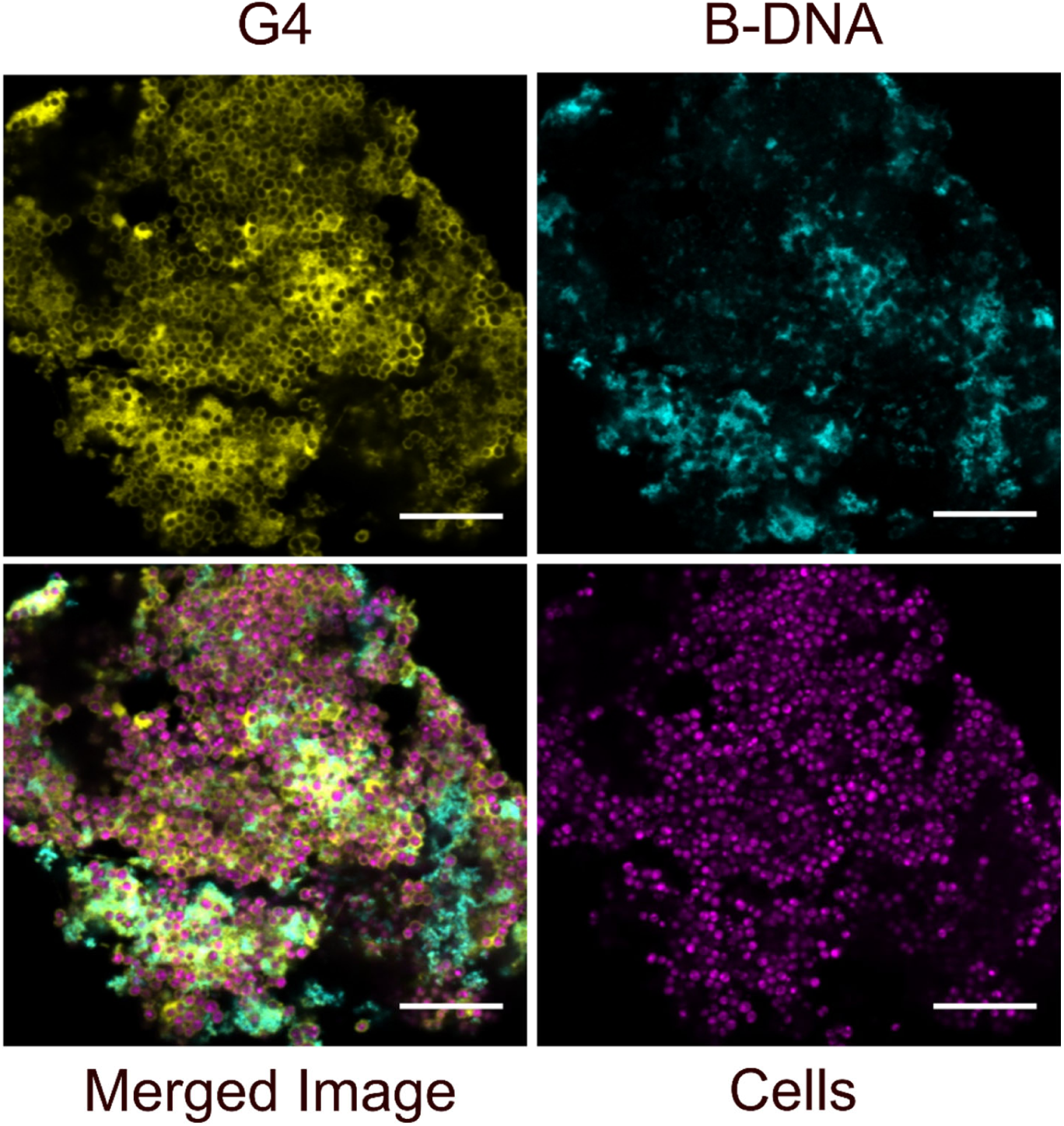
B-DNA only partially overlaps with the position of G4. Representative microscopy image of 10-day old biofilm showing G4 structures and B-DNA with regions of no-overlap after immunolabelling. B-DNA was not fully co-localised or binding to same G4 regions in the biofilm, indicating that the G4-specific antibody was binding to non-DNA structures. This implies that the G4 structures observed are a mixture of RNA and DNA.

**Extended Data Fig. 2.**
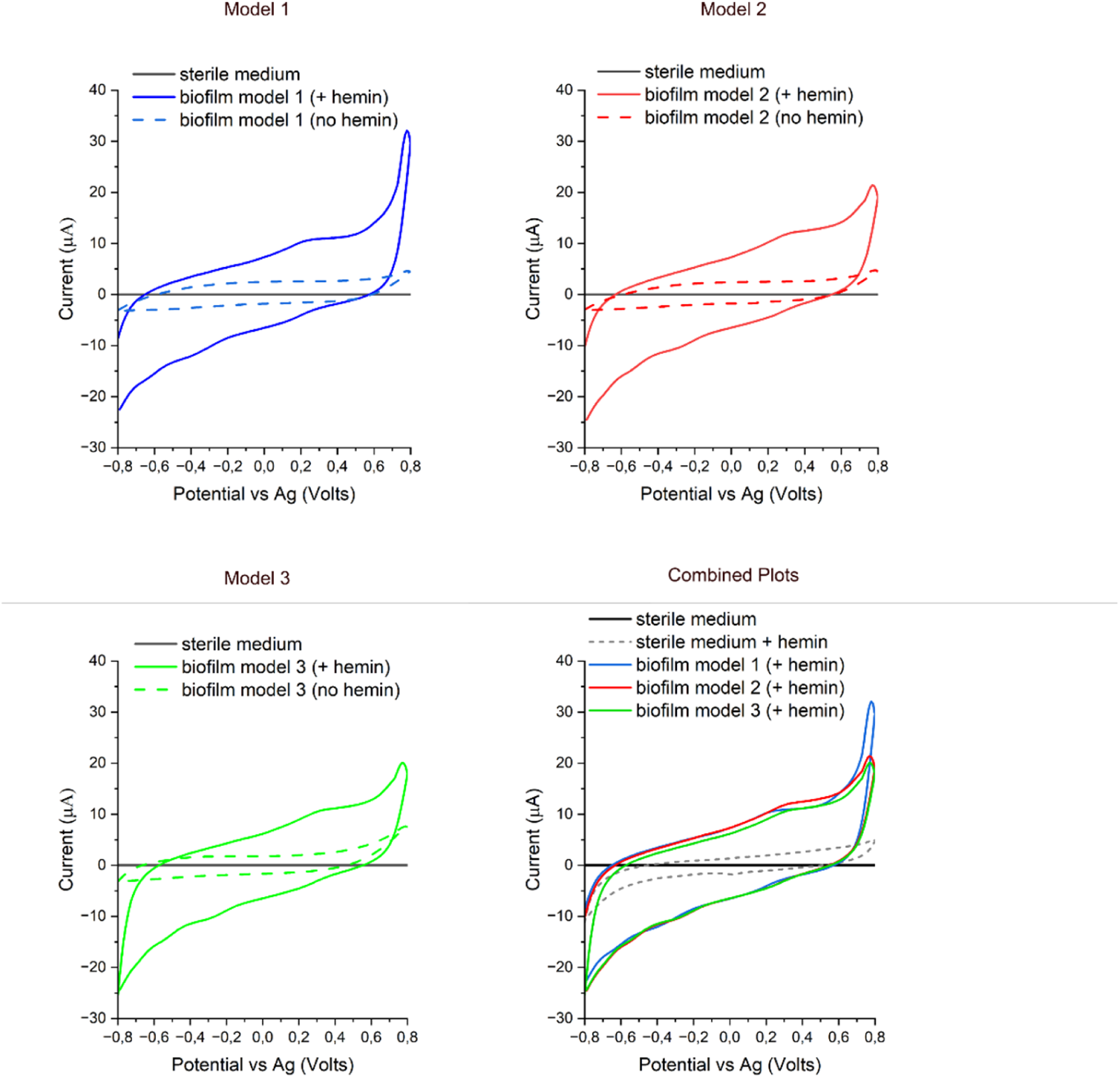
Cyclic Voltammogram (CV) of hemin (10 µM) treated, and untreated biofilms of model 1, model 2 and model 3 formed by *S. aureus*. All measurements were obtained after 72-hour culture of pre-grown biofilms which had been grown for 4 d (model 1), 8 d (model 2) and 12 d (model 3). Sterile BHI medium (supplemented with 0.2 M NaCl), BHI medium (+0.2 M NaCl and +10 µM hemin), and biofilms grown in BHI/NaCl medium without hemin supplementation served as controls.

**Extended Data Fig. 3.**
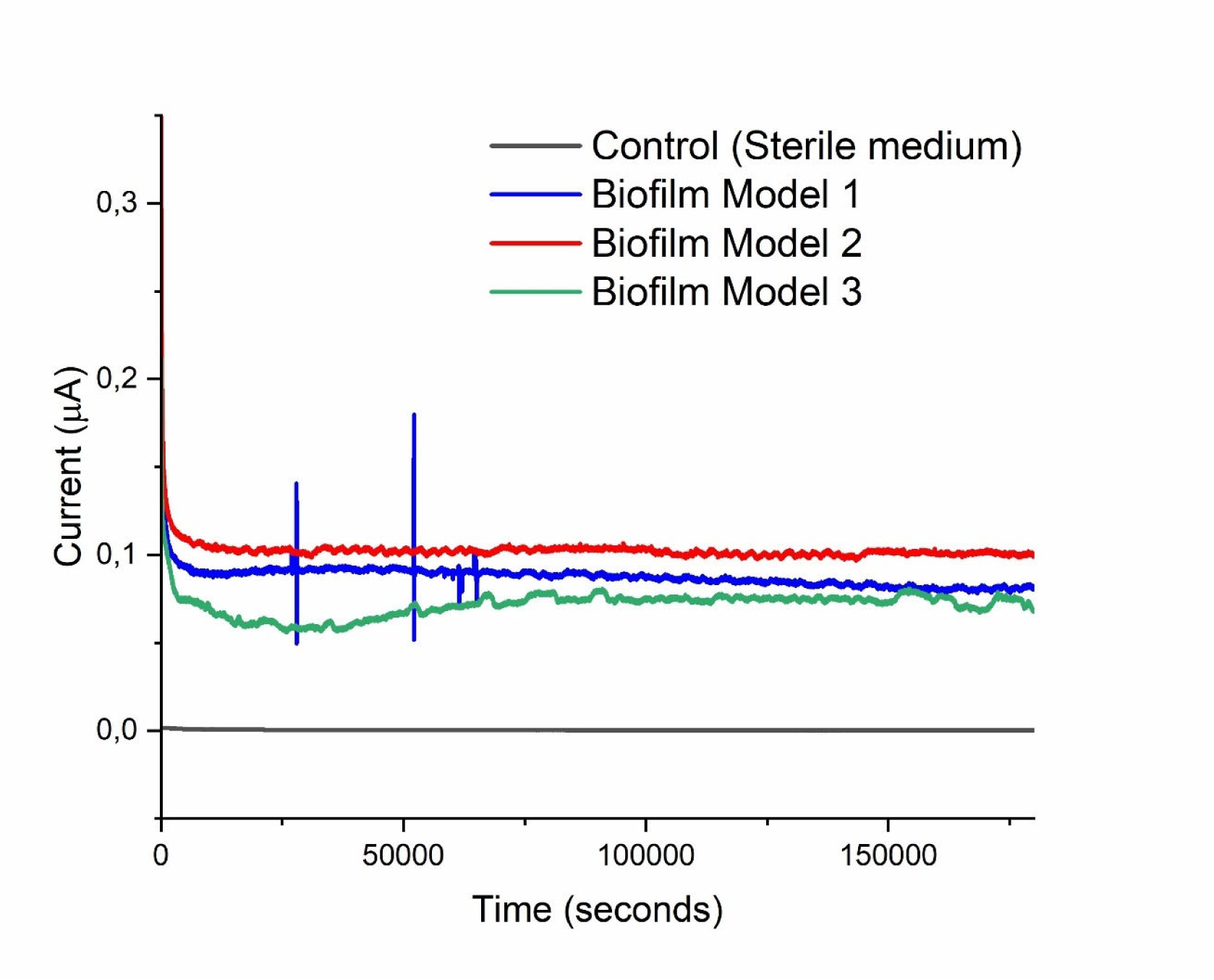
Chronoamperometric traces show stable electric current generation from all three electroactive biofilm models. Aerobically formed biofilms of models 1, 2 and 3, were transferred to anaerobic conditions with addition of fresh medium and hemin (10 µM) and adaptation on electrodes. Electroactive adaptation was at poised potential of +400 mV for 72 h to stabilize electron flow in biofilm. Current production from stabilised electroactive biofilms were then measured for extra 180,000 s (50 h). Chronoamperometric measurements were recorded after electroactive growth for 72 h for biofilms and extra 50 h measurements show long-term stability of electroactivity in biofilm models.

**Extended Data Fig. 4.**
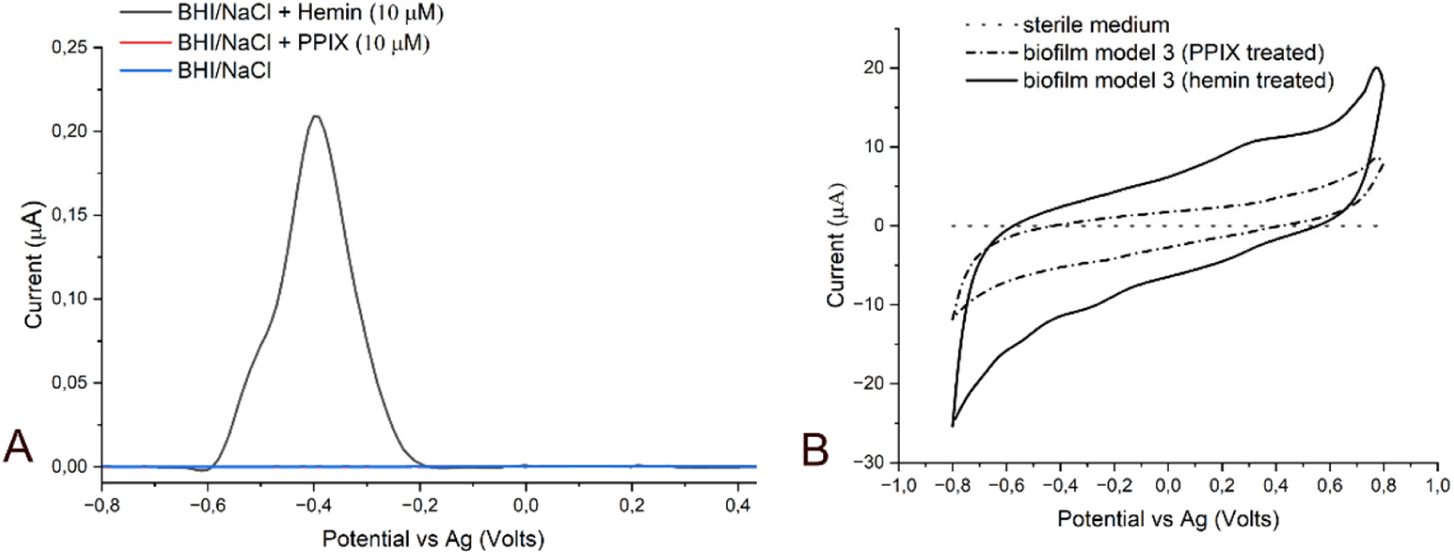
Representative voltammograms showing influence of medium supplementation with hemin and protoporphyrin IX (PPIX) on electrochemical properties of both medium and biofilms grown in the medium. (A) Differential pulse voltammogram (DPV) of sterile BHI/NaCl medium supplemented with hemin in comparison with sterile BHI/NaCl medium supplemented with protoporphyrin IX (PPIX). Both hemin and PPIX were at 10 µM concentrations (B) Representative cyclic voltammogram (CV) of hemin and PPIX treated medium treated biofilms formed by *S. aureus* biofilm model 3. Both treatments of hemin and PPIX were at the concentration of 10 µM, and measurements were obtained after 72h anaerobic culture of pre-grown biofilms of model 2. Sterile BHI supplemented with 0.2 M NaCl Medium served as control.

**Extended Data Fig. 5.**
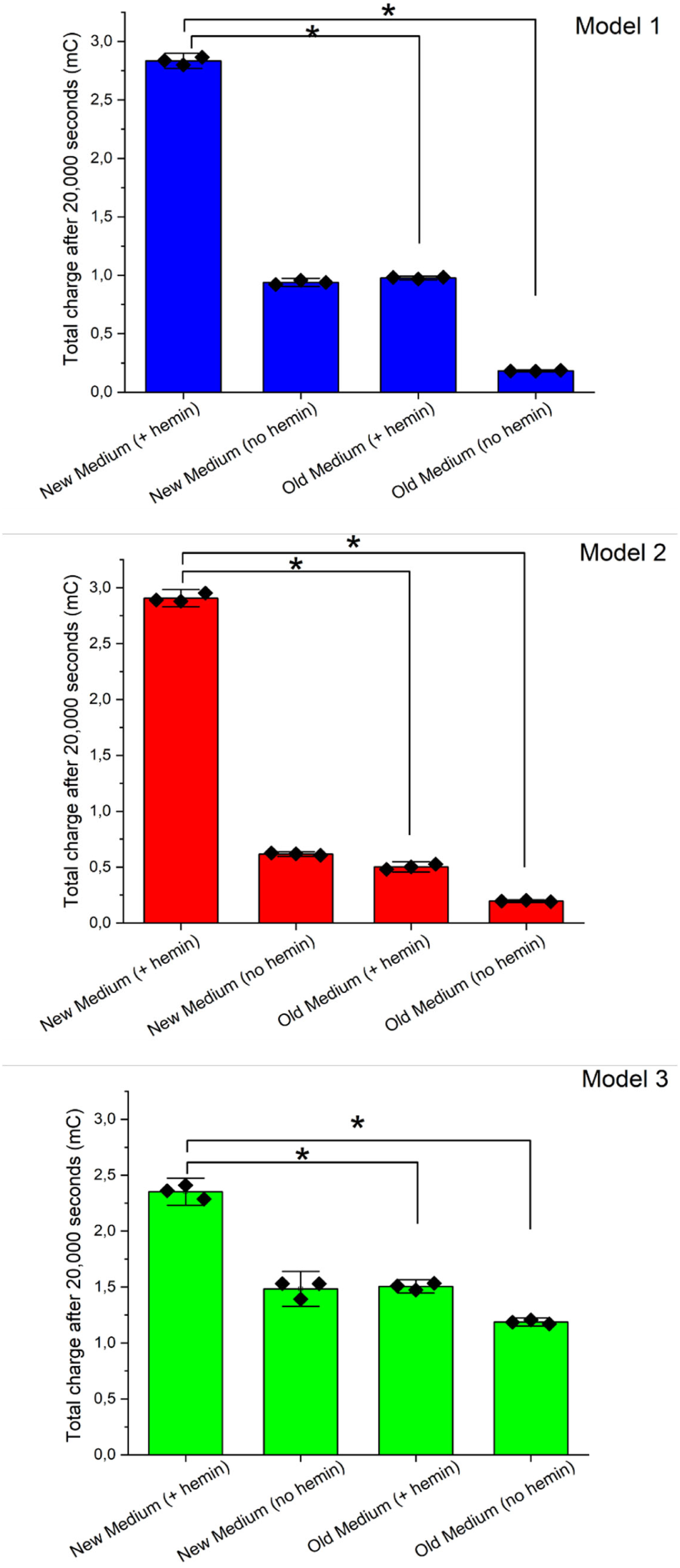
Comparative electron flow effects of hemin addition and nutrient replenishment on biofilm models after transfer of aerobically grown biofilms to anaerobic electroactive conditions. Charge measurements were done after 20,000 s (5.6 h) of addition of hemin and/fresh medium with +400 mV poised potential. A combination of hemin and fresh medium when transferred to anaerobic conditions generated optimum electron flow in all biofilm models. Old medium (with and without hemin) and new medium (without hemin) had low impacts on electron flow. Asterisks above the bars indicate significant differences (*p < 0.05).

## Notes

### Competing Interest Statement

The authors have declared no competing interest.

